# Protein Retrieval via Integrative Molecular Ensembles (PRIME) through extended similarity indices

**DOI:** 10.1101/2024.03.19.585783

**Authors:** Lexin Chen, Arup Mondal, Alberto Perez, Ramón Alain Miranda-Quintana

**Affiliations:** Department of Chemistry, University of Florida, FL, USA; Quantum Theory Project, University of Florida, FL, USA

**Keywords:** algorithms, protein structure, molecular simulation, clustering, ensembles

## Abstract

Molecular dynamics (MD) simulations are ideally suited to describe conformational ensembles of biomolecules such as proteins and nucleic acids. Microsecond-long simulations are now routine, facilitated by the emergence of graphical processing units. Processing such ensembles on the basis of statistical mechanics can bring insights about different biologically relevant states, their representative structures, states, and even dynamics between states. Clustering, which groups objects based on structural similarity, is typically used to process ensembles, leading to different states, their populations, and the identification of representative structures. For some purposes, such as in protein structure prediction, we are interested in identifying the representative structure that is more similar to the native state of the protein. The traditional pipeline combines hierarchical clustering for clustering and selecting the cluster centroid as representative of the cluster. However, even when the first cluster represents the native basin, the centroid can be several angstroms away in RMSD from the native state – and many other structures inside this cluster could be better choices of representative structures, reducing the need for protein structure refinement. In this study, we developed a module—Protein Retrieval via Integrative Molecular Ensemble (PRIME), that consists of tools to determine the most prevalent states in an ensemble using extended continuous similarity. PRIME is integrated with our Molecular Dynamics Analysis with *N* -ary Clustering Ensembles (MDANCE) package and can be used as a post-processing tool for arbitrary clustering algorithms, compatible with several MD suites. PRIME was validated with ensembles of different protein and protein complex systems for their ability to reliably identify the most native-like state, which we compare to their experimental structure, and to the traditional approach. Systems were chosen to represent different degrees of difficulty such as folding processes and binding which require large conformational changes. PRIME predictions produced structures that when aligned to the experimental structure were better superposed (lower RMSD). A further benefit of PRIME is its linear scaling – rather than the traditional O(*N* ^2^) traditionally associated to comparisons of elements in a set.

## Introduction

Proteins are dynamic systems, with closely coupled structure/function relationships. Some proteins fold into well-defined structures that help us to identify their functional role and mechanism of action. In many cases, these mechanisms involve accessing different biologically relevant structural states, which might happen on time or length scales inaccessible to experiments. By following picosecond to picosecond changes, molecular simulations produce ensembles that bring insights about different states accessible to the system. In some scenarios, such simulations start from an experimentally determined state, and conventional MD sampling extensively can access nearby biologically relevant states.^1–3^ In these cases we typically want to identify each sub-state, and their relative importance. In other scenarios, the native structure might be unknown but can be found through enhanced sampling approaches. For example, in protein structure determination starting from extended states or in the study of complexes starting from a known receptor and ligand situated far apart. In this scenario, users are typically interested in identifying the most native-like state predicted by the system. Clustering helps in both cases – in the first by identifying the boundaries by closely related sub-states, their relative importance (based on cluster population, and their representative structures. In the second scenario, we are typically concerned with the ability to identify native-like states, which is ideally represented in the highest population cluster – due to limitations in force fields and sampling strategies, this state is not always sampled in simulations, and when sampled, it is not necessarily the highest population cluster.

Ensembles produced by MD samples from a Boltzmann distribution should be processed using the principles of statistical mechanics. Clustering has been an appealing tool using structural similarity to distinguish between different states in the ensemble and provide information about their relative importance based on populations. It removes the ”cherrypicking” of structures from the ensemble by systematic ways of reporting representative structures as the centroid of a cluster.^4,5^

Various methodologies exist to segment data into meaningful groups, but the question remains, how do you get the best representative structure. However, the cluster definition might itself affect what the cluster centroid is, and thus the quality of the prediction. While this is often not very relevant when starting from native states, it can have a significant impact on structure prediction. As an example, the Critical Assessment of Structure Prediction event was started thirty years ago to describe the state of the art of the field in predicting the structures of proteins given their sequence. Given the tight deadlines and the number of targets, MD approaches are not very suitable for this competition, where homology modeling, statistical potentials,^6–8^ and more recently machine learning (e.g. Alphafold^9^ and RoseTTAFold^10^) have had greater success. Nonetheless, it is a perfect scenario for blind testing of the force field and sampling capacities of MD-based approaches. ^11,12^ In this scenario, predicting tight or dispersed clusters will result in representative structures that are very different from each other and the experimental structure.

Traditionally, hierarchical clustering^13^ has been used for determining clusters and cluster centroids. The primary obstacle lies in obtaining ensembles that encompass the native protein structure. Once such an ensemble is acquired, pinpointing the most significant centroid presents another significant challenge. However, this is a common problem for any clustering method. The most popular approach to this problem is to either use the centroid or medoid of the most populated cluster as a proxy for the native structure. However, as can be seen in Fig. 1, even if we obtain clusters containing the native structure, still selecting the right representative structure is a challenging problem. Oftentimes, this leads to estimates removed from the experimental structure in more than 3 *Å*. For example, any structure to the left of the red line in Fig. 1 would have been a better candidate, as quantified by the RMSD separation to the native conformation.

**Figure 1:**
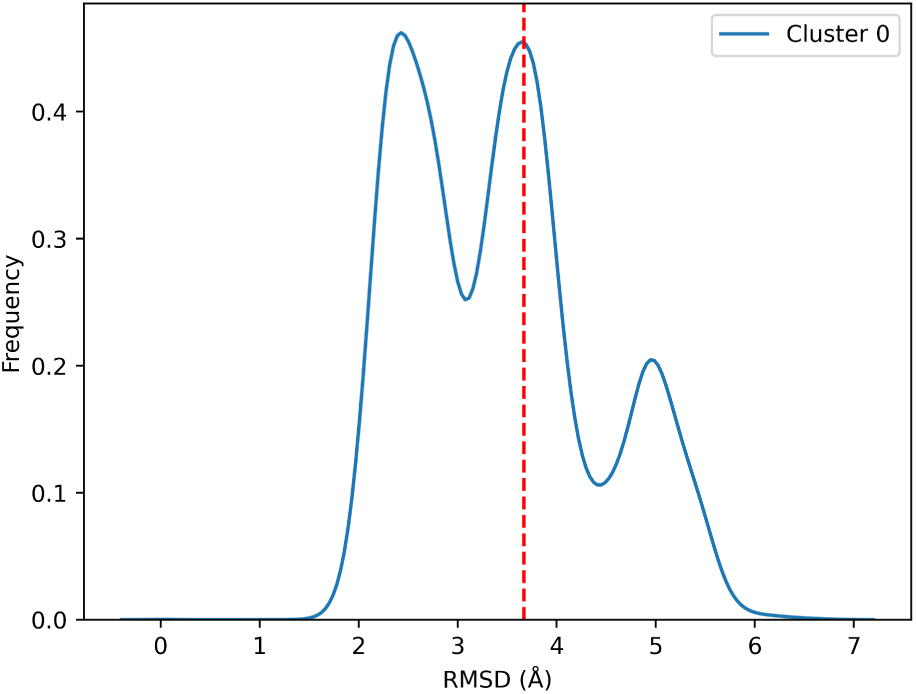
The RMSD distribution of the top (most populated) cluster referenced to the native structure for the system, low-temperature REMD N0968s1. The red line represents the RMSD of the medoid to the experimental native structure.

**Figure 2:**
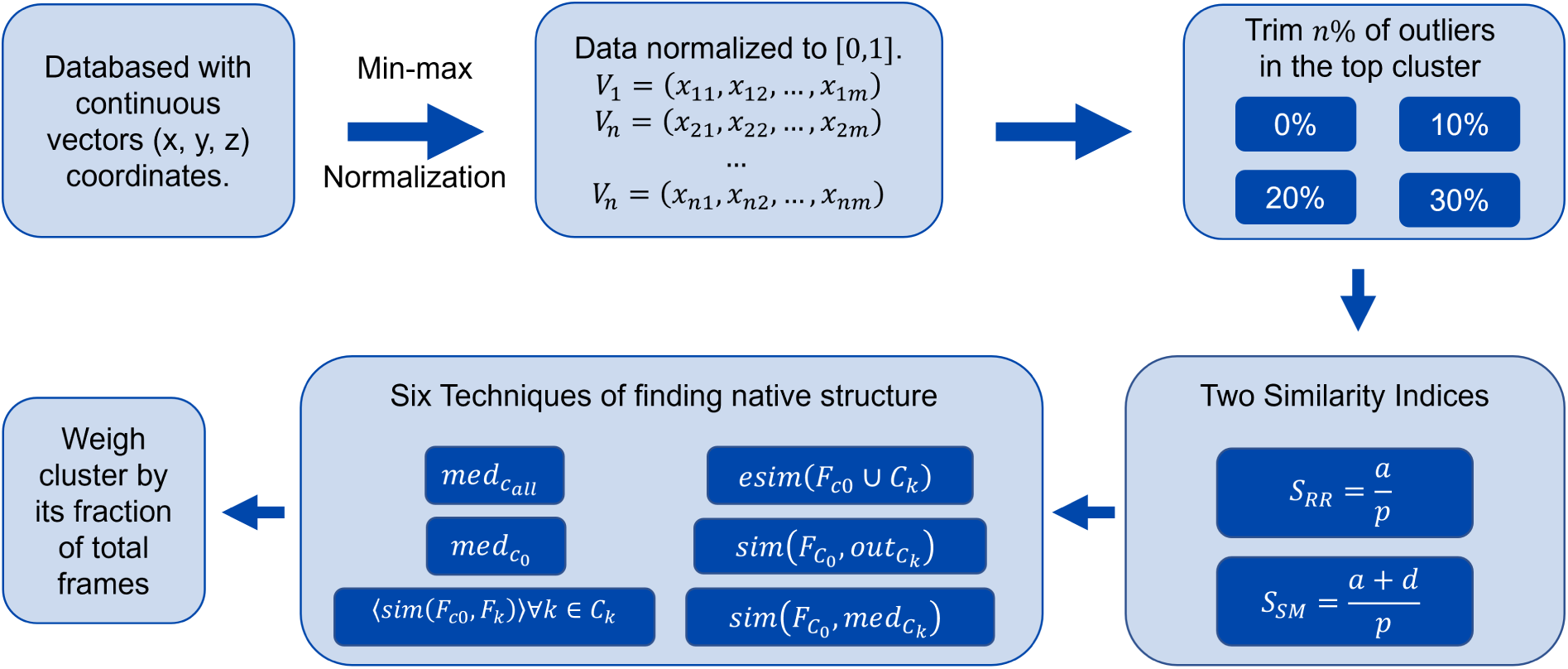
Flow chart of PRIME implementation.

**Figure 3:**
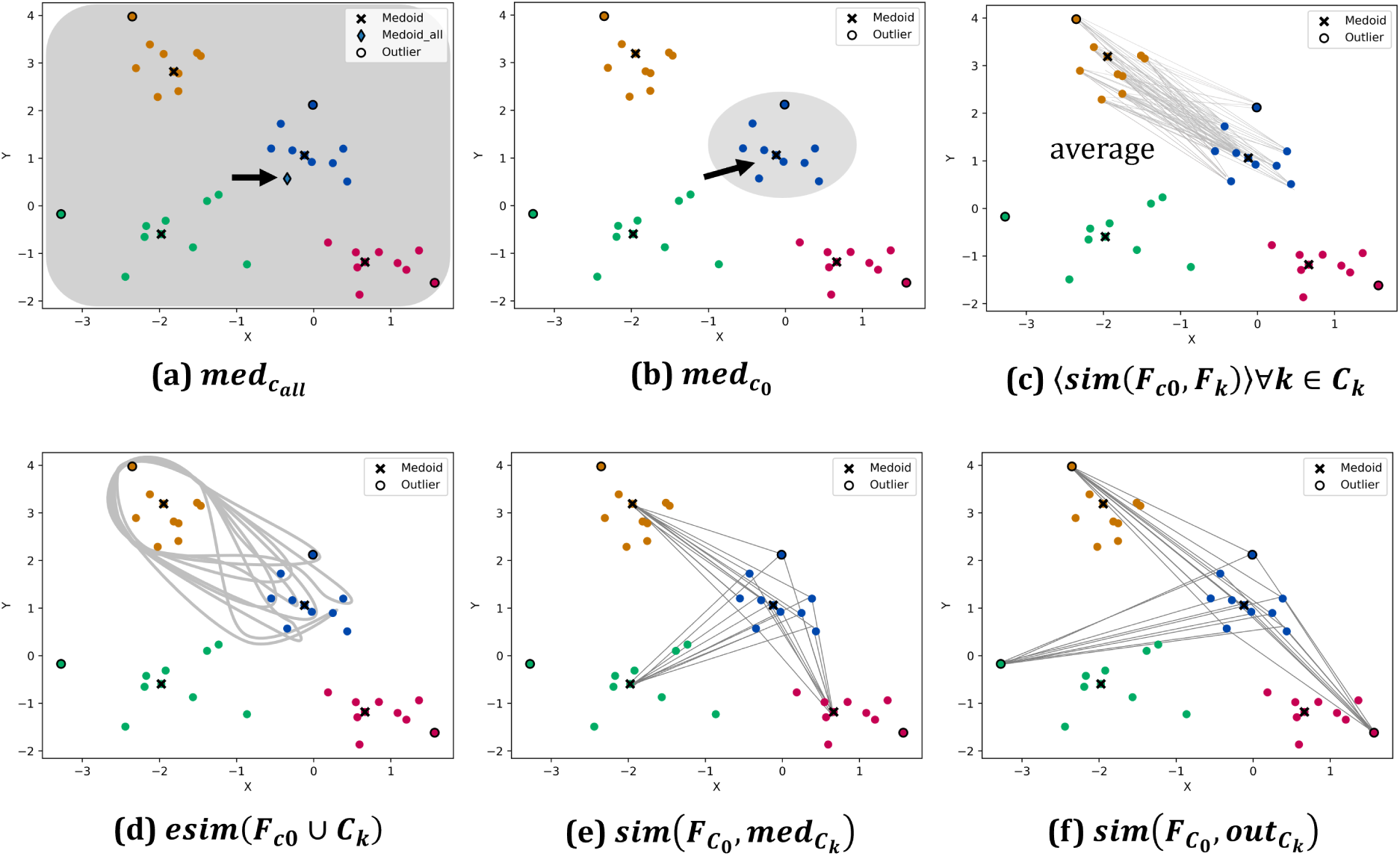
Six methods of native structure determination. All plots are 2D sample data and each color represents a different cluster. The blue cluster is the *c*_0_, the most populated cluster.

Here we propose to leverage the computational efficiency of extended similarity indices, which can compare multiple compounds in linear scaling across various datasets, ^14–19^ to try to identify better cluster representatives. Identifying the medoid within a dataset has been a challenge in computer science because contemporary algorithms operate with time complexity of *O*(*N* ^2^) or resort to approximations to achieve a *O*(*N* log *N* ) time complexity.^20^ We have found a way to solve this problem in *O*(*N* ) using extended similarity. The key is the concept of complementary similarity,^21^ which is also instrumental for the implementation of new structure retrieval protocols. We developed a package—Protein Retrieval via Integrative Molecular Ensembles (PRIME), that consists of tools to help identify closely related conformations using the extended continuous similarity. PRIME was able to map all the structural motifs in the studied systems and required unprecedented linear scaling.

In this work, we first introduce the protein structure prediction using extended similarity and then apply different metrics to several systems including protein folding ensembles and protein-peptide interaction ensembles. Our results show that our methods using PRIME were able to retrieve representative structures from the ensemble in better agreement with the native structure than the standard centroid method. We have made the code freely available at https://github.com/mqcomplab/PRIME.

## Theory

### Hierarchical Agglomerative Clustering

Hierarchical agglomerative clustering (HAC) initially assigns each data point as a cluster and then merges similar clusters at every iteration. Various merging techniques are commonly employed: single, centroid, and average linkages. Single linkage merges clusters based on the distance between the closest points of two clusters. On the other hand, complete linkage merges clusters based on the maximum distance between any two points in the clusters. Centroid linkage merges clusters based on the minimum distance between the centroids of two clusters Average linkage merges clusters based on the minimum average distance between all pairs of points from the two clusters. Among these techniques, average linkage and centroid linkage are preferred because they tend to produce compact and capable of identifying well-separated clusters of different shapes and sizes.^22^ Single linkage, although it offers computational efficiency, frequently leads to the formation of elongated clusters,^23^ whereas complete linkage tends to produce smaller clusters. HAC suffers from limitations such as time complexity of *O*(*N* ^2^) due to the need to compute pairwise similarities at each iteration,^24,25^ sensitivity to outliers depending on the linkage method used,^22^ and dependency on data permutation when deciding how to merge clusters with identical similarities.^26^ However, despite the power and versatility of HAC, the way that we use this clustering (or any other clustering algorithm for what matters), to identify native structures is rather simplistic. That is, one just identifies the most populated cluster and then selects the centroid of that basin. This effectively ignores all the information from the remaining clusters. In the rest of this contribution, we see how we could incorporate information from the other clusters to improve the prediction of native conformations.

### Extended Similarity

Extended similarity is a linear scaling approach of comparing the similarity of *n* objects or observations at the same time. The extended continuous similarity is an extension of the extended similarity, in which its input is an *N* × *D* matrix. *N* is the number of frames or observations and *D* is the spatial coordinates of each frame.^17,19^ To ensure the matrix aligns with extended similarity, it is necessary to normalize it within [0, 1], to match the scale of cheminformatics metrics. This involves scaling the values between the minimum and maximum coordinates, thus maintaining consistency with ranking from RMSD calculation. This 2D matrix will be transformed into a 1D vector by summing the features column-wise across all samples. This vector can then be utilized to compute the extended similarity, a global measure indicating the similarity of a set. Conversely, applying the complementary similarity enables a local approach or the ranking of objects to extract information like the medoid or outlier.

Using complementary similarity, we can identify the medoid of a set in linear scaling. That is, given a set of conformations, the complementary similarity of any element will be the extended similarity of all the conformations, except the chosen one. So, the element with the lowest complementary similarity will correspond to the medoid of the set. We will use this to rank conformations, from the medoid to the outlier. This will be the key to implementing new structure retrieval protocols.

### PRIME

To identify the native structure of a system, several preprocessing steps are necessary. Starting with a 2D matrix, it is normalized to [0, 1] to accommodate for cheminformatics indices (More on this step in Extended Similarity Section). Since there may be some ”noise” in the data, trimming the outliers in the top (most populated) cluster will lead to more robust results. Therefore, it undergoes four options of trimming to prune the outliers from the top clusters to better identify the high-density region. For the first time, trimming is possible in *O*(*N* ) due to the complementary similarity to efficiently rank the most representative to the most outlier-like. Subsequently, two similarity indices were investigated to determine which similarity indices were used for the six techniques of finding native structures to identify the most native-like structure better, Russel-Rao (RR) and Sockal-Michener (SM).

Then, six different techniques were tested to find the native structure. These six techniques aim to efficiently establish relations between sub-sets of conformations (as opposed to the traditional single-cluster strategy), thus pinpointing better representative structures. Lastly, clusters are weighted by the ratio of the cluster population to the total population since less uniformly distributed clusters should not have a lowly populated cluster to bias the native structure calculation.

A common technique for native structure prediction has been using clustering to find an ensemble with a native-like structure and then find the centroid of the top cluster. The extended similarity protein retrieval framework includes six ways of structure prediction, including outlier trim, all with linear scaling. There are six ways of identifying more relevant protein structures in the ensemble. After clustering, the structure is typically determined from the most populous cluster, disregarding information from *n-1* clusters. PRIME offers an alternative for refining the quality of predicted structures with the integration of additional data.

1. *med_c_*_0_ , where *c*_0_ is the top (most populated) cluster. (2) *med_c__all_* , where *med* is the medoid and *c_all_*is all clusters. (3) ⟨*sim*(*F_c_*_0_ *, F_k_*)⟩∀*k* ∈ *C_k_*, where *sim* is the pairwise similarity between two objects; *F_c_*_0_ is a frame from the top cluster; *F_k_* is a frame from cluster *k* ; *C_k_*is cluster *k*. (4) *esim*(*F_c_*_0_ ∪ *C_k_*), where *esim* is the extended similarity between the objects. (4) *sim*(*F_c_*_0_ *, med_C__k_* ). (5) *sim*(*F_c_*_0_ *, out_C__k_* ), where *out* is outlier. These six techniques consider the relationship between the clusters found to predict the native structure of a protein with various advantages and disadvantages.

*med_c_*_0_ calculates the medoid of the top cluster after clustering using complementary similarity. After clustering, in this case, HAC, similar structures are grouped and the most populated cluster is also the most dominant structure in the simulation. By pinpointing the native structure to a smaller range, this technique allows for an efficient and simple solution to predict the protein structure in complex simulations. Therefore, *med_c_*_0_ is a popular method in the community for protein structure prediction and the *med_c_*_0_ untrimmed will be the benchmarking method for other methods in PRIME. Although it can serve as an adequate estimate of the native structure, it can fail when cluster sizes are not uniformly distributed, or the cluster shape is arbitrary or non-convex. Henceforth, PRIME is a potential solution to retrieve protein conformations of all clusters found and *med_c_*_0_ will be benchmarked against the other methods.

*med_c__all_* calculates the medoid of the whole dataset using complementary similarity. This method is the most efficient among the six techniques because it doesn’t necessitate any preliminary clustering, unlike the other five techniques. However, there is no way to weigh frame by importance. Therefore the medoid structure will most likely be located in the global minima, where the native structure is. However, in enhanced sampling MD simulation, some biasing factor is employed to favor the simulation to climb over the energy barriers; for example, in Replica Exchanged Molecular Dynamics (REMD), increasing the temperature will facilitate jumping over energy barriers between basins.

⟨*sim*(*F_c_*_0_ *, F_k_*)⟩∀*k* ∈ *C_k_*—*pairwise* method—involves calculating the average pairwise similarity between a frame in the top cluster and every frame in cluster *k*. This process is repeated for each frame in the top cluster and then repeated for every cluster. In other words, this technique is a condensed pairwise matrix between a subset of the data (top cluster) and all other clusters. Because the native structure is targeted to be in the top cluster, this method can pinpoint which frame in the top cluster is most dissimilar to all other structures in other clusters and the frame most similar to the frames in other clusters should be the native structure. In this case, weighing clusters by their population proved to be essential because more stable structures, usually found in more populated clusters, would be more represented and similar to the native structure than unstable structures by the energy barriers. The *pairwise* method is hypothesized to identify the protein structure closest to the native structure because it considers every observation in the clustering result. However, this advantage comes with a significant computational overhead as it scales *O*(*N* ^2^). *esim*(*F_c_*_0_ ∪*C_k_*)—*esim* method—entails computing the extended similarity between a frame in the top cluster and all frames in cluster *k*. This process is repeated for each frame in the top cluster and then repeated for each cluster. This method is very similar to the *pairwise* method above because it finds the similarity between a subset of data (top cluster) and other clusters. On the other hand, it does not utilize the pairwise similarity metrics but instead the extended similarity. Using the extended similarity, can efficiently speed up the calculation by condensing the 2D matrix into a single vector, the column sum of all vectors in the matrix. Thereby, it significantly speeds up the postprocessing analysis at *O*(*N* ) and still preserves similar results to the *pairwise* method.

*sim*(*F_c_*_0_ *, med_C__k_* )—*medoid-top* comparison—encompasses calculating the pairwise similarity between a frame in the top cluster and the medoid of cluster *k* (frame with the lowest complementary similarity). This procedure iterates for every frame within the top cluster and subsequently repeats for each cluster. From the *O*(*N* ) determination of the medoid of a subset, it can efficiently identify the medoid from a set. Compared to the *esim* method, this technique has further condensed the calculation from all observations to just the medoid. Theoretically, the *medoid-top* comparison may be more robust than the *esim* method because it does not consider outliers in the clusters. In contrast, the *esim* method considers all observations in a cluster *k* with no biasing factor for a particular observation. Therefore, the *medoid-top* comparison may be closer to the structure prediction the similarity between the medoid to the outlier of a cluster is significantly low or spherical. However, the *esim* method may be more accurate for clusters that are more non-convex and arbitrary because in these cases, the inclusion of all members would give a more accurate overview. This is also an *O*(*N* ) time complexity and can efficiently determine the protein structure.

*sim*(*F_c_*_0_ *, out_C__k_* )—*outlier-top* comparison—entails computing the pairwise similarity between a frame in the top cluster and the outlier of cluster *k* (frame with the highest complementary similarity). This procedure iterates for every frame within the top cluster and subsequently repeats for each cluster. The inclusion of this method can give an outlier of the compact of the cluster. Since both *medoid-top* and *outlier-top* comparisons are in place, the uniformity of the cluster can be inferred. With a similar result in both the medoid-top and *outlier-top*, this would infer there is great uniformity within the cluster and that the similarity between the medoid and the outlier is relatively high. The inclusion of this method can give insights into the quality of the clustering and see if this method is also a valid structure prediction technique.

## Methods

### Simulation Details

#### Protein folding ensembles based on the CASP13 NMR dataset

We used 15 protein sequences and unassigned NMR data provided in the NMR data assisted category of the 13^th^ edition of the Critical Assessment of Protein Structure Prediction (CASP13, https://predictioncenter.org/casp13/).^27^ These proteins range in size from 80 residues to 326 residues. We ran the MELD (Modeling Employing Limited Data)^28,29^ Bayesian inference approach incorporating the provided ambiguous NMR data (NOESY restraint type of data representing an unassigned spectrum of sparsely labeled proteins) data as distance restraints.^30^ Simulations were run for 0.5 *µ*s using Hamiltonian and Temperature Replica Exchange Molecular Dynamics (H, T-REMD) with 30 replicas. Protein target N0981D3 was extended to 1 *µ*s and N1005 to 1.5 *µ*s. We created two datasets from the ensembles at different temperatures (ranging from 300K at the lowest replica index to 550K at higher replicas). In the first dataset, we selected the ensembles corresponding to replicas at the lowest 5 temperatures – the *low-temperature data set*. The second dataset corresponds to replicas with a temperature range between 400K to 550K – the *high-temperature data set*, expected to sample a more diverse range of structures. Simulations were run using the gbNeck2 implicit solvent^31^ model along with the FF14SBside force field for proteins.^32,33^ We excluded target N0989 from the test sets due to its poor sampling. We analyzed the ensembles using a hierarchical clustering algorithm (with an epsilon cutoff of 2*Å*) from the CPPTRAJ package of AMBER suite^34^ to cluster the last half of the 5 lowest temperature replicas. The RMSDs of the representative structure (centroid of the top populated cluster) with respect to the native were used as the benchmark for this test set in this work.

#### The NMR Exchange Format dataset

We used 15 targets provided by the NMR Exchange Format (NEF) initiative (https://github.com/NMRExchangeFormat/NEF). These are small proteins, ranging from 76 to 202 residues, where the majority have highly flexible coil regions that confound typical clustering approaches. These targets have been solved by NMR and the NOESY data was also used in the predictions. This dataset is not sparsely labeled and thus much more data-rich than the previous datasets. We again used the MELD approach to combine the information and physics model in the generation of the ensembles using 30 replicas (with a temperature range between 300K and 550K changing in the first twelve replicas and remaining replicas simulated at 550K). Local distance restraints, resulting from peaks coming from neighboring residues (residue pairs separated by less than four residues) were enforced from the 1st replicas to the 24^th^ replicas with maximum strength i.e. 350 *kJmol^−^*^1^*nm^−^*^2^, and it was geometrically weakened to 0 at the 30^th^ replicas. The global distance restraints (peaks associated with residue pairs separated by four or more residues) were enforced from the 1st replicas to 12^th^ replicas (350 *kJmol^−^*^1^*nm^−^*^2^), then geometrically reduced to 0 at the 24^th^ replicas and no global distance restraints were enforced from the 24^th^ to 30^th^ replicas. This setup was used to facilitate the formation of the local contacts at the higher temperature replicas and then the global contacts as it came down the replica ladder. Each simulation was run for 50 ns with 100% of the NOESY peaks enforced. The FF14SBside force field and gbNeck2 implicit solvent model were again used. We used the five lowest temperature replicas to cluster using a hierarchical algorithm (with an epsilon cutoff of 1*Å*) based on RMSD of the residues with well-defined secondary structure. These representative structures were used as the third dataset.

#### Protein-Peptide complexes

Unlike previous test sets, the current systems entail the binding of intrinsically disordered peptides to a protein receptor.^35^ Here we used the ExtraTerminal (ET) domain of Bromodomain and ExtraTerminal domain (BET) family proteins as the receptors and we performed binding of six intrinsically disordered peptides that interact to ET domain of BETs with NMR assisted MELD simulations.^35^ We set up the simulations starting with the unbound structure of the receptor and an extended chain conformation for the peptide. In all cases, we used 3 types of NMR data: Chemical Shift Perturbation (CSP) data of the receptor, TALOS data derived from CSPs of the peptides, and three NOE peaks. CSP data was used to derive ambiguous distance restraints between the receptor and the peptides, TALOS data was used to restrain the backbone dihedrals of the peptide, and finally, more informative NOE peaks were used as unambiguous distance restraints. We transferred the CSP data of the ET-TP complex to other systems. Experimental TALOS and NOE data were used for ET-TP and ET-NSD3 whereas for JMJD6, LANA, CHD4, and BRG1 calculated TALOS and NOE data were used. All systems were simulated with 30 replicas for 1.5 *µ*s with the FF14SBside forcefield and gbNeck2 implicit solvent model. At the end of the simulations, the five lowest temperature replicas were extracted and the ensemble containing their last 1 *µ*s were used for clustering analysis with the previously described hierarchical clustering algorithm. We used an epsilon cutoff of 1.5Å for the peptide residues after aligning on the protein receptor. We ignored the flexible tail of the peptides from the clustering analysis (see our previous work for more details). ^35^ The RMSD of representative complexes to the experimental structures of the complex were used as the final dataset for the current study.

### Software Module

PRIME is released as a software module under Molecular Dynamics Analysis with *N* -ary Clustering Ensembles (MDANCE) package. The MDANCE GitHub repository can be found here: https://github.com/mqcomplab/MDANCE. PRIME provides postprocessing tools for analyzing clustering results that can be provided from MDANCE or other clustering platforms. It contains different trimming or similarity indices options, along with the six structural prediction methods for user modulation. MDANCE is a flexible and versatile clustering package that can perform various clustering, alignment, and seed selector methods and can interface with programs like AMBER^36^ and MDAnalysis.^37,38^ The PRIME GitHub repository can be found here: https://github.com/mqcomplab/PRIME.

## Results and discussion

The six techniques underwent validation with ensembles produced using different approaches and incorporating systems to represent different degrees of flexibility and common challenges such as folding or binding.

From Fig. 4-7, Eqn. 1 will determine how accurate each method is to the experimental native structure compared to the benchmarking method, *med_c_*_0_ . If the value is negative, then the method used agrees with the experimental native structure more than the bench-marking method. The bar graph indicates the fraction of systems with a negative value. A higher fraction would suggest a greater generality of this method. The superposition shows the overlaps between the best structure from PRIME methods to the experimental native structure.

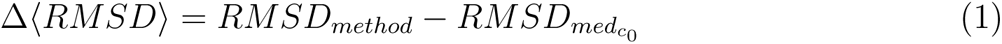

**Figure 4:**
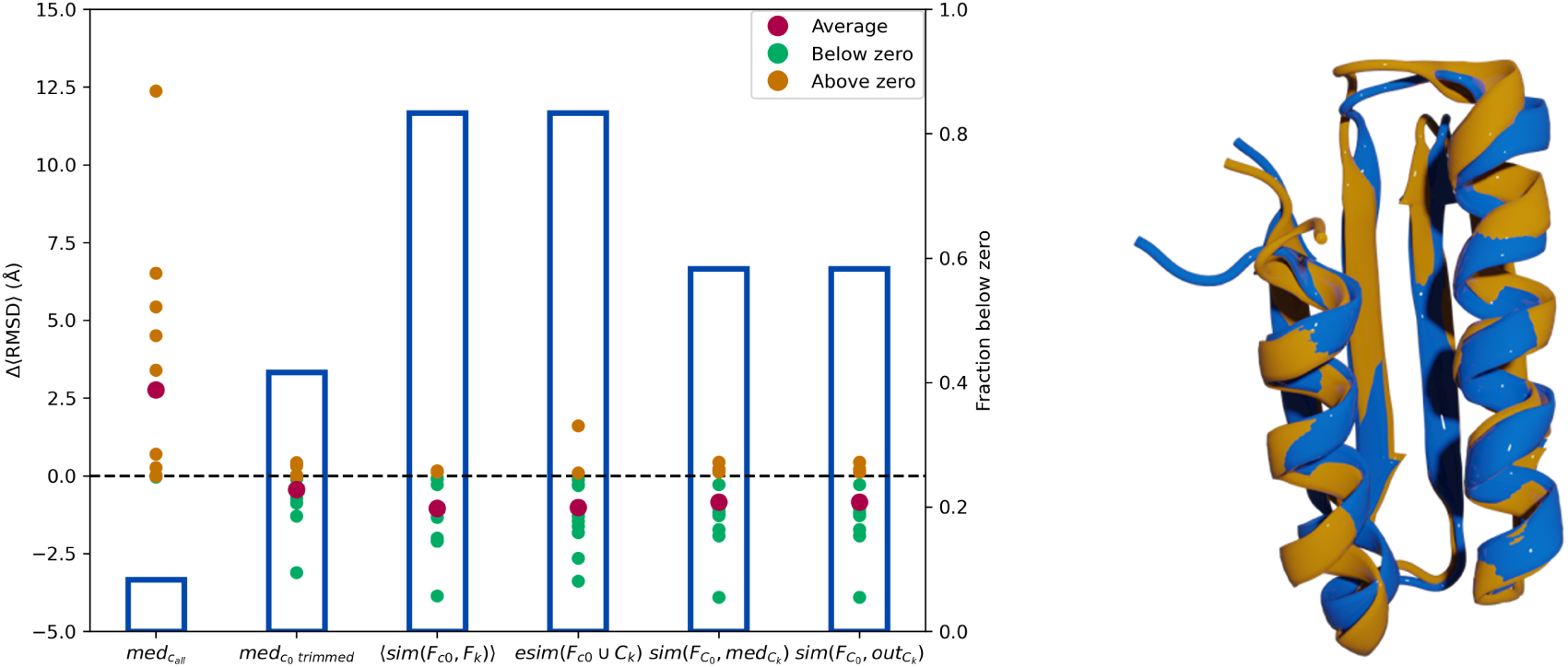
Left is Δ⟨*RMSD*⟩ of 12 low-temperature REMD systems between the native structure calculated with *med_c_*_0_ to the native structure calculated using other PRIME techniques. The scatter plot corresponds to the left y-axis and the bars correspond to the right y-axis. Right is a superposition of the most representative structures found with PRIME (yellow) and experimental native structures (blue) of n1008.

**Figure 7:**
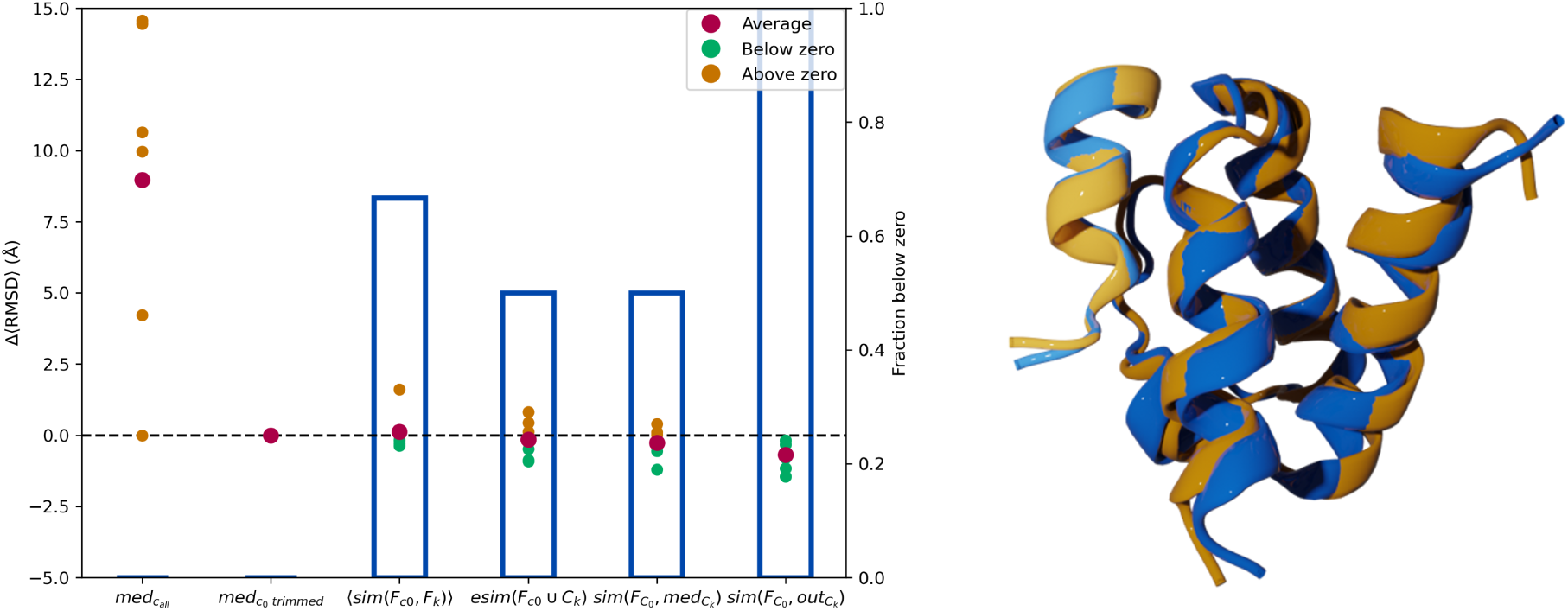
Left is Δ⟨*RMSD*⟩ of six protein-peptide REMD systems between native structure calculated with *med_c_*_0_ to native structure calculated using other PRIME techniques. The scatter plot corresponds to the left y-axis and the bars correspond to the right y-axis. Right is a superposition of the most representative structures found with PRIME (yellow) and experimental native structures (blue) of JMJD6.

In the low-temperature REMD systems, the *esim* method had the same fraction of systems falling below zero indicating a great generality in *esim* that is comparable to the more computationally costly pairwise method. The *medoid-top* method was able to significantly improve on the one benchmark method by more than 3.9 *Å*. Fig. 4 uses the SM index and a 10% trim. There is not much difference between the native structure calculated by *medoid-top* versus *outlier-top*. This can be due to the SM similarity index, which is not as susceptible to detecting outliers as the RR index. Because RR puts more emphasis on the presence of a shared feature, whereas SM has equal weight on the presence or absence of a shared feature, RR can more effectively capture distinctive features. However, SM can more effectively predict protein structure, which can be because SM is a metric but RR is non-metric.^39^ Since it is a metric, it more resembles the nature of RMSD, which is a distance metric.

Analysis of high-temperature ensembles exhibits a drop in performance, most likely associated with the higher conformational diversity observed in these conditions. Fig. 5 uses the SM index and a 10% trim. Since it is a higher temperature it might not spend as much time in the native structure as in lower temperatures.^40^ Here, the *esim* method has a higher generality than all other methods in recovering better representative structures, even the pairwise method. However, it can also be noted that the *medoid-top* and *outlier-top* have one system that is significantly greater than zero. The *medoid-top* comparison usually struggles with more arbitrary cluster shapes because it typically works best when clusters are spherical. Therefore, the *esim* method might produce a more accurate protein prediction in these cases. Remarkably, the performance of the *esim* method is roughly the same in the low and high-temperature regimes. This highlights the generality of the *n*-ary similarity measures, given by their ability to capture multiple correlations simultaneously.

**Figure 5:**
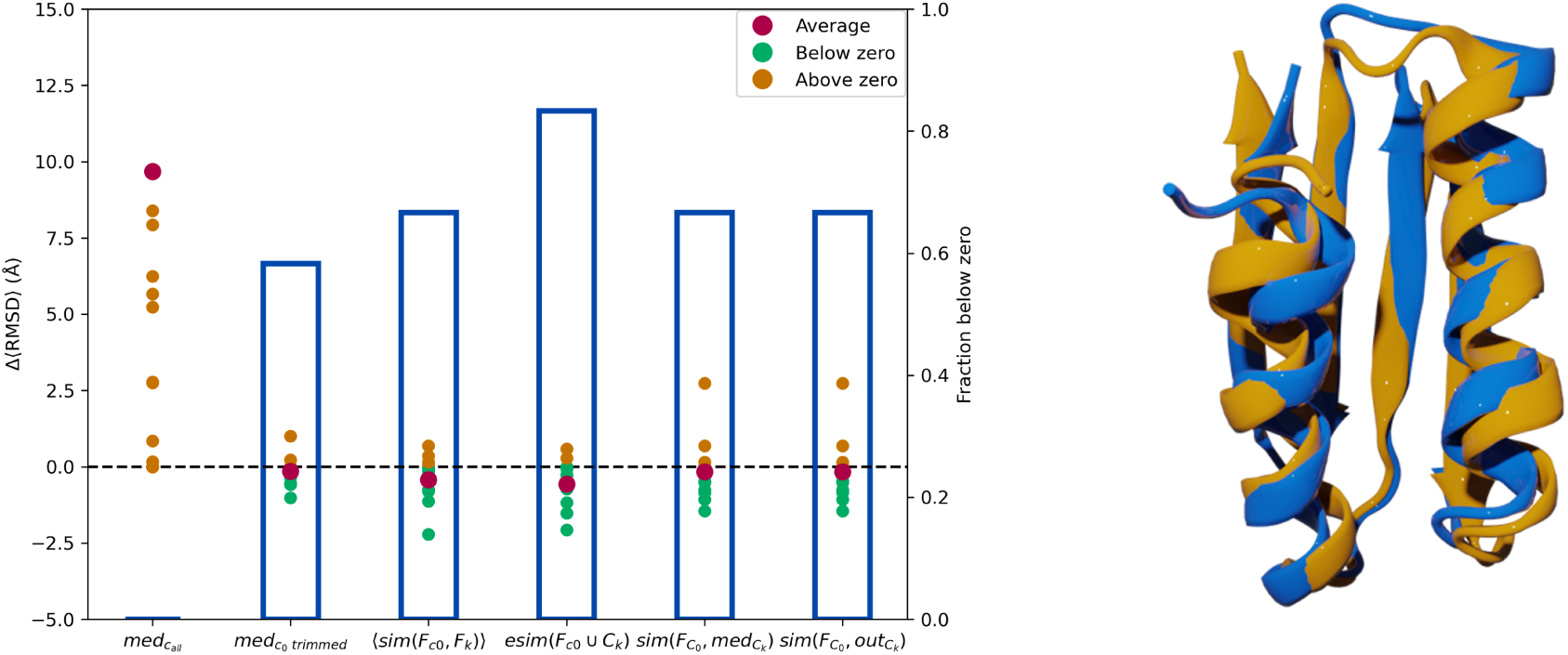
Left is Δ⟨*RMSD*⟩ of twelve high-temperature REMD systems between native structure calculated with *med_c_*_0_ to native structure calculated using other PRIME techniques. The scatter plot corresponds to the left y-axis and the bars correspond to the right y-axis. Right is a superposition of the most representative structures found with PRIME (yellow) and experimental native structures (blue) of n1008.

The fifteen flexible-protein systems are more complicated than the two different temperature REMD systems because there are many flexible loops/motifs and contain more residues. Due to the REMD, it can explore many configurations. Fig. 6 uses the SM index and a 30% trim. Since these protein systems are very flexible, a decrease in the generality and accuracy can be observed. Once again, the *esim* method holds up substantially with more than 60% of the time better than the benchmarking method. What is even more noteworthy, even without a perfect agreement in the absolute coordinate values, the *esim* method was able to match even the most flexible local patterns in the native conformation.

**Figure 6:**
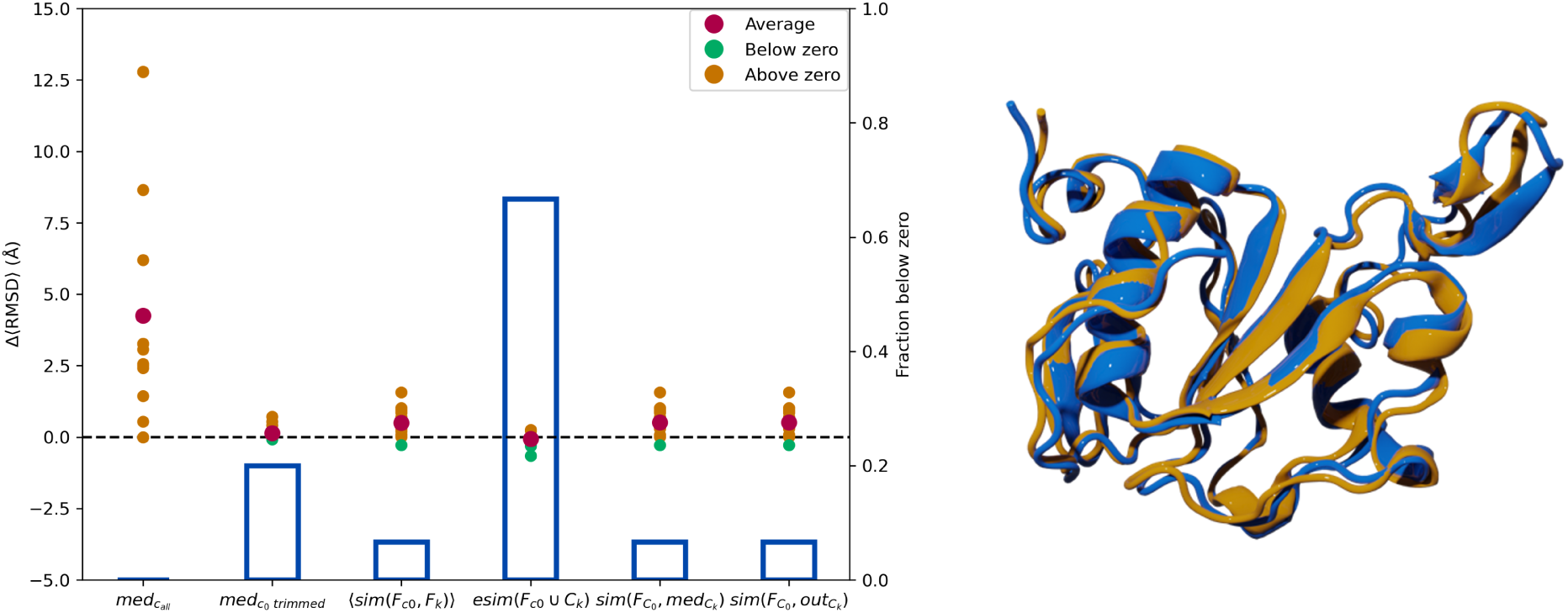
Left is Δ⟨*RMSD*⟩ of fifteen flexible-protein REMD systems between native structure calculated with *med_c_*_0_ to native structure calculated using other PRIME techniques. The scatter plot corresponds to the left y-axis and the bars correspond to the right y-axis. Right is a superposition of the most representative structures found with PRIME (yellow) and experimental native structures (blue) of 2k2e.

Lastly, in the protein-peptide systems, a 100% generality can be observed with the *outlier-top* method. Fig. 7 uses the RR index and a 20% trim. The *outlier-top* was able to predict accurately for all the systems compared to the benchmarking method. From the overlap, there is a very close agreement between the peptide structure predicted to bind to the protein and the actual native structure. In this case, the *esim* method severely underperformed when compared to the *outlier-top* comparison when it comes to the fraction of predictions that were better than the benchmark results. However, for the systems that were corrected by the *esim* method, the magnitude of this improvement was similar to those seen with the *outlier-top* comparison.

Overall, the esim method consistently improves upon the traditional approach of focusing only on the medoid of the top cluster, in the cases when there is only one molecule in the simulation. On average, the improvement for the low-T, high-T, and flexible protein systems was -3.38*Å*, -2.07*Å*, and -0.66*Å*. Remarkably, the average gains for the low-T and high-T simulations were comparable, which speaks of the generality of the method. On the other hand, as noted above, the two-molecule systems were better described by the comparison to the outliers (average improvement of -1.76*Å* A).

## Conclusion

We have shown that there is virtue in rethinking how we choose representative structures from clustered data. As an example, we have taken the application to predict the most native-like structure of a biological system. In these conditions, ensembles are typically clustered, and then the centroid of the top cluster is chosen to represent the native state. The performance of the whole method is typically assessed by how far the centroid is from the native state – leaving out better representative structures in the top cluster. This leads to the refinement of such predictions, typically with more detailed MD analysis.

Our new metrics present an attractive alternative since we can improve upon the single-cluster strategy, at no additional computational cost. That is, we obtain structures that are, on average, -1.89 *Å* better, and in some individual cases, this improvement can even reach -3.90 *Å*. The *esim* method consistently outperformed the other variants for the low-T, high-T, and flexible protein cases, while the outlier variant improved all the peptide-protein systems. In either case, the underlying motivation is the same: we can use information from the least populated clusters to augment our knowledge of the most populated one.

The (dismal) performance of the *med_c__all_* approach (just finding the medoid of all the frames) highlights the key importance of clustering in refining conformations. In this regard, PRIME presents a key advantage, since it can be used after any clustering or data segmentation step. Moreover, PRIME only requires a list of cluster labels, so there is no need to generate or store the expensive *O*(*N* ^2^) matrix of pairwise RMSD between conformations. This is possible due to the *n*-ary similarity metrics, which allow comparing the frames in the most populated cluster to the entirety (or a subset) of the other clusters with unmatched efficiency.

## Acknowledgement

RAMQ and LC thank support from the National Institute of General Medical Sciences of the National Institutes of Health under award number R35GM150620. AP and AM thank NIH-NIGMS under award number R01GM149646.

## References

(1) Rickey Welch, G.; Somogyi, B.; Damjanovich, S. The role of protein fluctuations in enzyme action: A review. Progress in Biophysics and Molecular Biology 1982, 39, 109–146.

(2) Wrabl, J. O.; Gu, J.; Liu, T.; Schrank, T. P.; Whitten, S. T.; Hilser, V. J. The role of protein conformational fluctuations in allostery, function, and evolution. Biophysical Chemistry 2011, 159, 129–141.

(3) Frauenfelder, H.; Chen, G.; Berendzen, J.; Fenimore, P. W.; Jansson, H.; McMahon, B. H.; Stroe, I. R.; Swenson, J.; Young, R. D. A unified model of protein dynamics. Proceedings of the National Academy of Sciences 2009, 106, 5129–5134.

(4) Shortle, D.; Simons, K. T.; Baker, D. Clustering of low-energy conformations near the native structures of small proteins. Proceedings of the National Academy of Sciences 1998, 95, 11158–11162.

(5) Harder, T.; Borg, M.; Boomsma, W.; Røgen, P.; Hamelryck, T. Fast large-scale clustering of protein structures using Gauss integrals. Bioinformatics 2012, 28, 510–515.

(6) Hiranuma, N.; Park, H.; Baek, M.; Anishchenko, I.; Dauparas, J.; Baker, D. Improved protein structure refinement guided by deep learning based accuracy estimation. Nature Communications 2021, 12, 1340.

(7) Ovchinnikov, S.; Park, H.; Kim, D. E.; DiMaio, F.; Baker, D. Protein structure prediction using Rosetta in CASP12. *Proteins: Structure*, Function, and Bioinformatics 2018, 86, 113–121.

(8) Park, H.; Lee, G. R.; Kim, D. E.; Anishchenko, I.; Cong, Q.; Baker, D. High-accuracy refinement using Rosetta in CASP13. Proteins: Structure, Function, and Bioinformatics 2019, 87, 1276–1282, Publisher: John Wiley & Sons, Ltd.

(9) Jumper, J. et al. Highly accurate protein structure prediction with AlphaFold. Nature 2021, 596, 583–589.

(10) Baek, M. et al. Accurate prediction of protein structures and interactions using a three-track neural network. 2021,

(11) Perez, A.; Morrone, J. A.; Brini, E.; MacCallum, J. L.; Dill, K. A. Blind protein structure prediction using accelerated free-energy simulations. Science Advances 2016, 2, e1601274, tex.pmcid: PMC5106196 tex.rating: 5.

(12) He, Y.; Mozolewska, M. A.; Krupa, P.; Sieradzan, A. K.; Wirecki, T. K.; Liwo, A.; Kachlishvili, K.; Rackovsky, S.; Jagiela, D.; Slusarz, R.; Czaplewski, C. R.; Oldziej, S.; Scheraga, H. A. Lessons from application of the UNRES force field to predictions of structures of CASP10 targets. Proceedings of the National Academy of Sciences of the United States of America 2013, 110, 14936 – 14941, tex.pmcid: PMC3773777 tex.rating: 0.

(13) Daura, X.; Gunsteren, W. F. v.; Mark, A. E. Folding–unfolding thermodynamics of a beta-heptapeptide from equilibrium simulations. Proteins: Structure, Function, and Bioinformatics 1999, 34, 269–280.

(14) Miranda-Quintana, R. A.; Bajusz, D.; Rácz, A.; Héberger, K. Extended similarity indices: the benefits of comparing more than two objects simultaneously. Part 1: Theory and characteristics†. Journal of Cheminformatics 2021, 13, 32.

(15) Miranda-Quintana, R. A.; Rácz, A.; Bajusz, D.; Héberger, K. Extended similarity indices: the benefits of comparing more than two objects simultaneously. Part 2: speed, consistency, diversity selection. Journal of Cheminformatics 2021, 13, 33.

(16) Miranda-Quintana, R. A.; Bajusz, D.; Rácz, A.; Héberger, K. Differential Consistency Analysis: Which Similarity Measures can be Applied in Drug Discovery? Molecular Informatics 2021, 40, 2060017.

(17) Rácz, A.; Dunn, T. B.; Bajusz, D.; Kim, T. D.; Miranda-Quintana, R. A.; Héberger, K. Extended continuous similarity indices: theory and application for QSAR descriptor selection. Journal of Computer-Aided Molecular Design 2022, 36, 157–173.

(18) Rácz, A.; Bajusz, D.; Héberger, K. Life beyond the Tanimoto coefficient: similarity measures for interaction fingerprints. Journal of Cheminformatics 2018, 10, 48.

(19) Rácz, A.; Mihalovits, L. M.; Bajusz, D.; Héberger, K.; Miranda-Quintana, R. A. Molecular Dynamics Simulations and Diversity Selection by Extended Continuous Similarity Indices. Journal of Chemical Information and Modeling 2022, 62, 3415–3425.

(20) Eppstein, D.; Wang, J. Fast Approximation of Centrality. Journal of Graph Algorithms and Applications 2004, 8, 39–45.

(21) Chang, L.; Perez, A.; Miranda-Quintana, R. A. Improving the analysis of biological ensembles through extended similarity measures. Physical Chemistry Chemical Physics 2022, 24, 444–451.

(22) Shao, J.; Tanner, S. W.; Thompson, N.; Cheatham, T. E. Clustering Molecular Dynamics Trajectories: 1. Characterizing the Performance of Different Clustering Algorithms. Journal of Chemical Theory and Computation 2007, 3, 2312–2334.

(23) Torda, A. E.; Van Gunsteren, W. F. Algorithms for clustering molecular dynamics configurations. Journal of Computational Chemistry 1994, 15, 1331–1340.

(24) Glielmo, A.; Husic, B. E.; Rodriguez, A.; Clementi, C.; Nóe, F.; Laio, A. Unsupervised Learning Methods for Molecular Simulation Data. Chemical Reviews 2021, 121, 9722– 9758.

(25) Xu, D.; Tian, Y. A Comprehensive Survey of Clustering Algorithms. Annals of Data Science 2015, 2, 165–193.

(26) Bakkelund, D. Order preserving hierarchical agglomerative clustering. Machine Learning 2022, 111, 1851–1901.

(27) Sala, D. et al. Protein structure prediction assisted with sparse NMR data in CASP13. *Proteins: Structure*, Function, and Bioinformatics 2019, 87, 1315–1332.

(28) MacCallum, J. L.; Perez, A.; Dill, K. Determining protein structures by combining semireliable data with atomistic physical models by Bayesian inference. Proceedings of the National Academy of Sciences 2015, 112, 6985 – 6990.

(29) Perez, A.; MacCallum, J. L.; Dill, K. A. Accelerating molecular simulations of proteins using Bayesian inference on weak information. Proceedings of the National Academy of Sciences of the United States of America 2015, 112, 11846–51.

(30) Mondal, A.; Perez, A. Simultaneous Assignment and Structure Determination of Proteins From Sparsely Labeled NMR Datasets. Frontiers in Molecular Biosciences 2021, 8, 774394.

(31) Nguyen, H.; Roe, D. R.; Simmerling, C. Improved Generalized Born Solvent Model Parameters for Protein Simulations. Journal of chemical theory and computation 2013, 9, 2020 – 2034.

(32) Maier, J. A.; Martinez, C.; Kasavajhala, K.; Wickstrom, L.; Hauser, K. E.; Simmerling, C. ff14SB: Improving the Accuracy of Protein Side Chain and Backbone Parameters from ff99SB. Journal of chemical theory and computation 2015, 11, 3696 – 3713.

(33) Hornak, V.; Abel, R.; Okur, A.; Strockbine, B.; Roitberg, A.; Simmerling, C. Comparison of multiple Amber force fields and development of improved protein backbone parameters. Proteins 2006, 65, 712 – 725.

(34) Roe, D. R.; Cheatham, T. E. PTRAJ and CPPTRAJ: Software for Processing and Analysis of Molecular Dynamics Trajectory Data. Journal of Chemical Theory and Computation 2013, 9, 3084–3095.

(35) Mondal, A.; Swapna, G.; Lopez, M. M.; Klang, L.; Hao, J.; Ma, L.; Roth, M. J.; Montelione, G. T.; Perez, A. Structure Determination of Challenging Protein–Peptide Complexes Combining NMR Chemical Shift Data and Molecular Dynamics Simulations. Journal of Chemical Information and Modeling 2023, 63, 2058–2072.

(36) Case, D. et al. *Amber* 2020 ; University of California, San Francisco, 2020.

(37) Gowers, R.; Linke, M.; Barnoud, J.; Reddy, T.; Melo, M.; Seyler, S.; Domański, J.; Dotson, D.; Buchoux, S.; Kenney, I.; Beckstein, O. MDAnalysis: A Python Package for the Rapid Analysis of Molecular Dynamics Simulations. Austin, Texas, 2016; pp 98–105.

(38) Michaud-Agrawal, N.; Denning, E. J.; Woolf, T. B.; Beckstein, O. MDAnalysis: A toolkit for the analysis of molecular dynamics simulations. Journal of Computational Chemistry 2011, 32, 2319–2327.

(39) Zhang, B.; Srihari, S. N. Binary vector dissimilarity measures for handwriting identification. Document recognition and retrieval X. 2003; pp 28 – 38.

(40) Stumpff-Kane, A. W.; Maksimiak, K.; Lee, M. S.; Feig, M. Sampling of near-native protein conformations during protein structure refinement using a coarse-grained model, normal modes, and molecular dynamics simulations. *Proteins: Structure*, Function, and Bioinformatics 2008, 70, 1345–1356.

