## Supplementary Information for "Protein Retrieval via Integrative Molecular Ensembles (PRIME) through extended similarity indices"

### Supporting Information Available

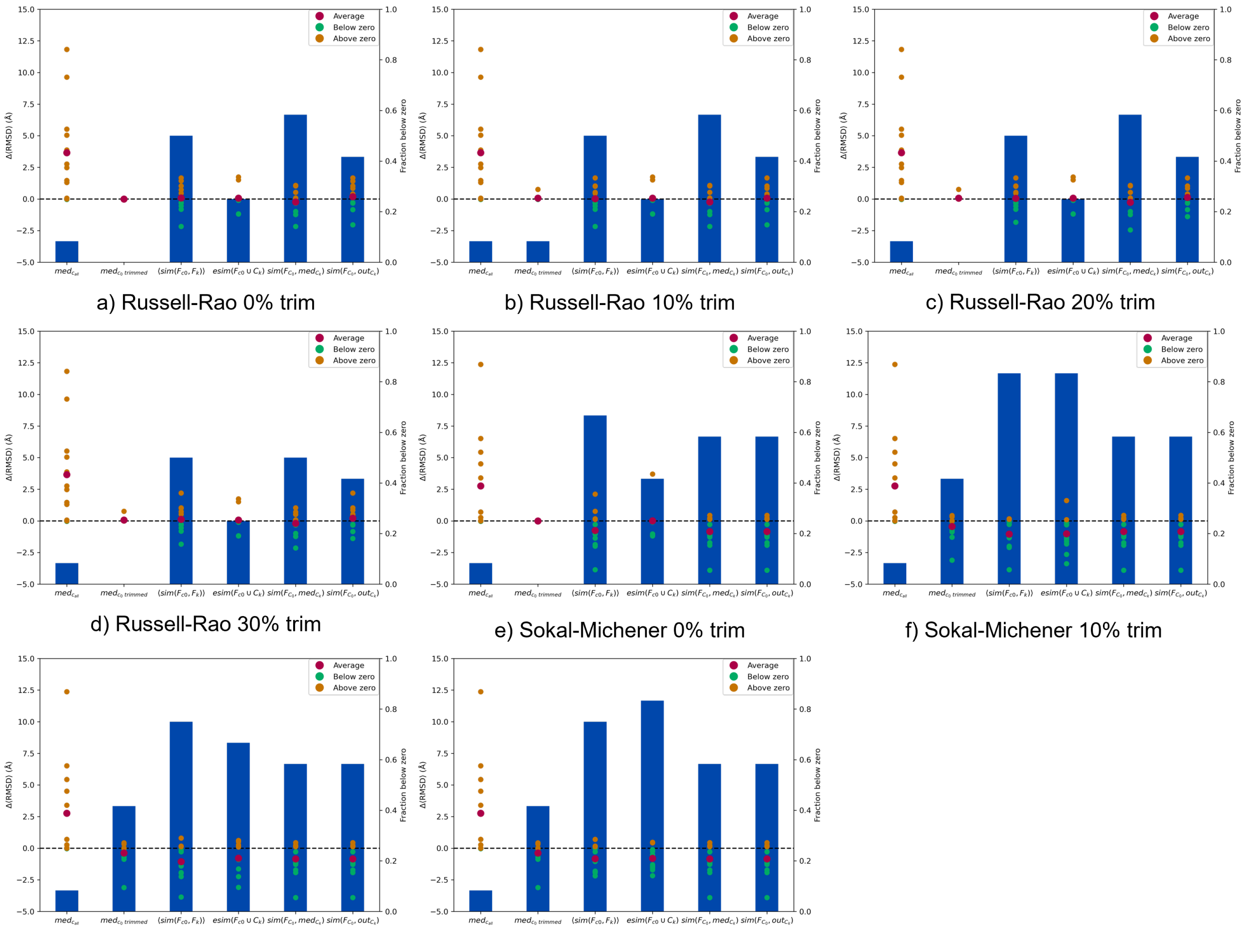


a) Russell-Rao 0% trim

b) Russell-Rao 10% trim

c) Russell-Rao 20% trim

d) Russell-Rao 30% trim

e) Sokal-Michener 0% trim

f) Sokal-Michener 10% trim

g) Sokal-Michener 20% trim

h) Sokal-Michener 30% trim

Figure S1: $\Delta\langle RMSD\rangle$ of twelve low-temperature REMD systems between native structure calculated with $med_{c0}$ to native structure calculated using other PRIME techniques. The scatter plot corresponds to the left y-axis and the bars correspond to the right y-axis. (a)-(h) uses the indicated similarity indices and trimming.

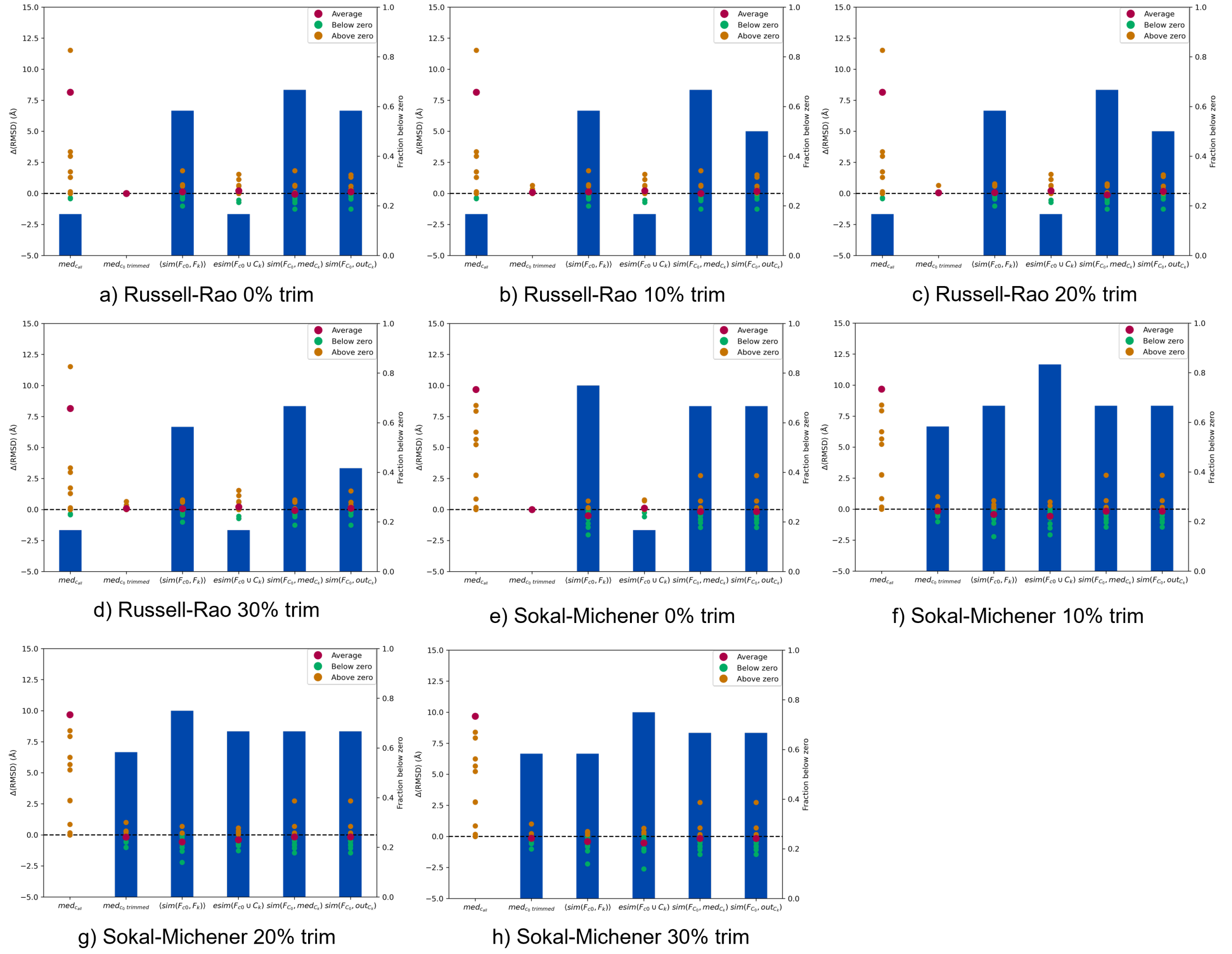


a) Russell-Rao 0% trim
b) Russell-Rao 10% trim
c) Russell-Rao 20% trim
d) Russell-Rao 30% trim
e) Sokal-Michener 0% trim
f) Sokal-Michener 10% trim
g) Sokal-Michener 20% trim
h) Sokal-Michener 30% trim

Figure S2: $\Delta\langle RMSD\rangle$ of twelve high-temperature REMD systems between native structure calculated with $med_{c_0}$ to native structure calculated using other PRIME techniques. The scatter plot corresponds to the left y-axis and the bars correspond to the right y-axis. (a)-(h) uses the indicated similarity indices and trimming.

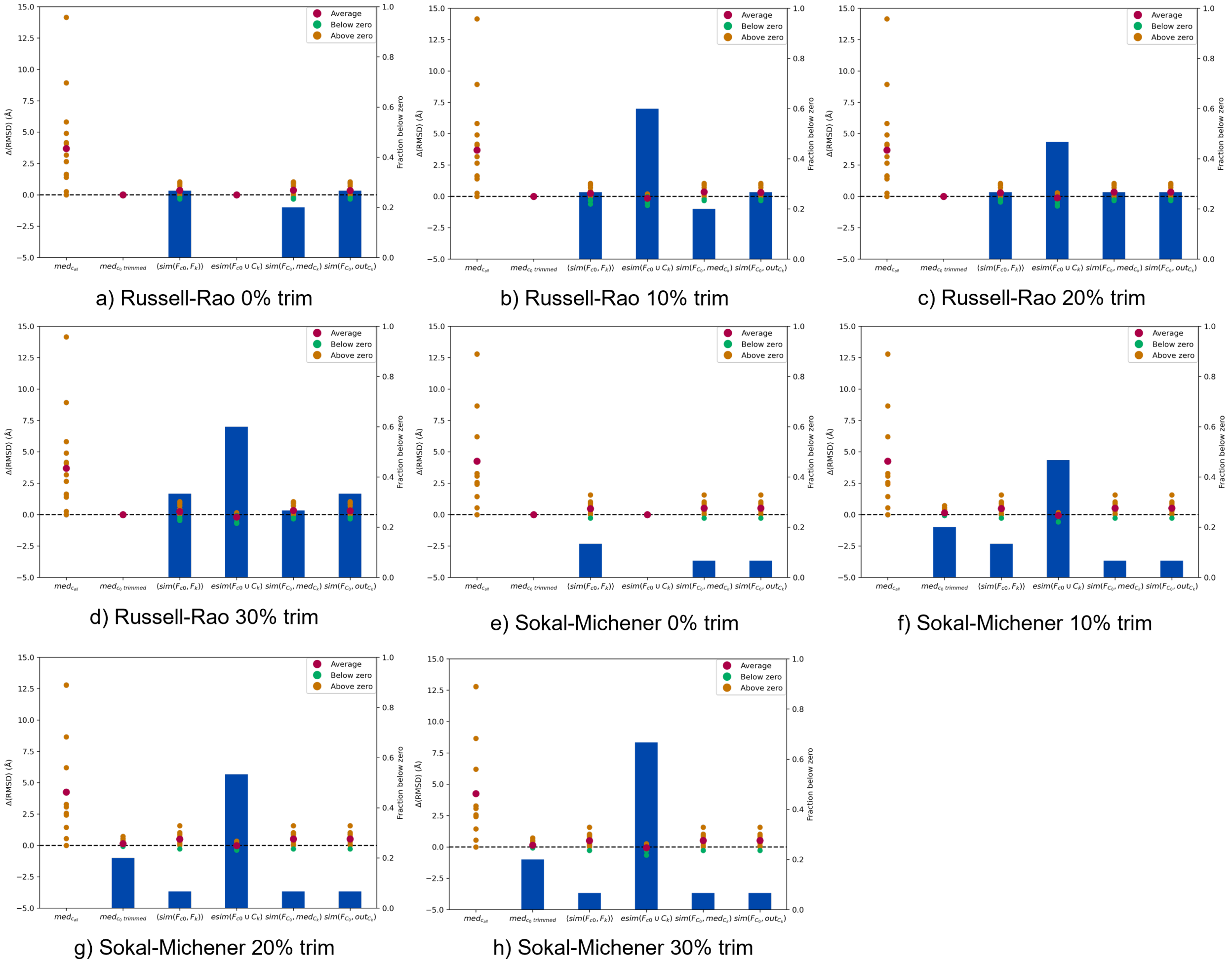


Figure S3: $\Delta\langle RMSD\rangle$ of fifteen flexible-protein REMD systems between native structure calculated with $med_{c0}$ to native structure calculated using other PRIME techniques. The scatter plot corresponds to the left y-axis and the bars correspond to the right y-axis. (a)-(h) uses the indicated similarity indices and trimming.

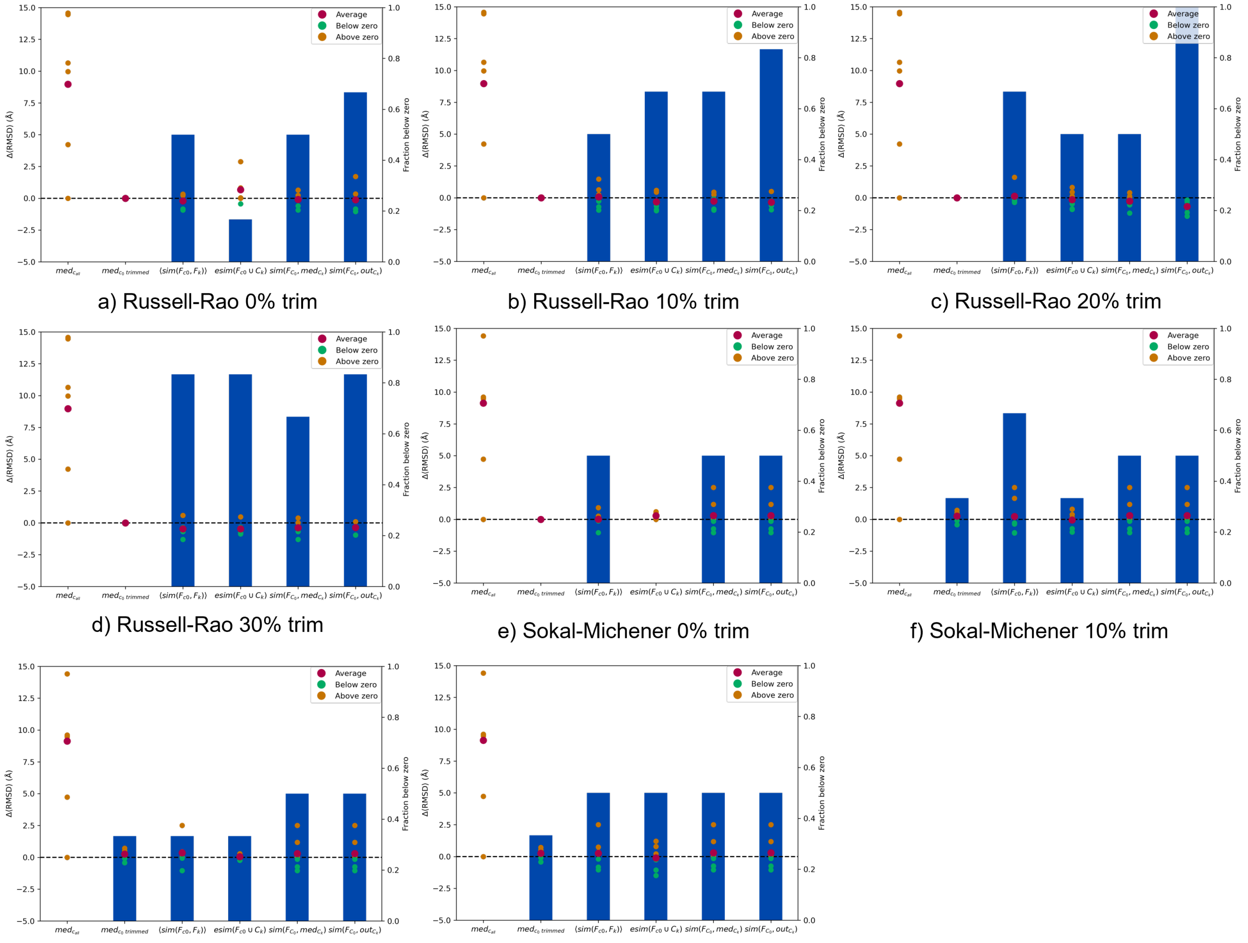


a) Russell-Rao 0% trim

b) Russell-Rao 10% trim

c) Russell-Rao 20% trim

d) Russell-Rao 30% trim

e) Sokal-Michener 0% trim

f) Sokal-Michener 10% trim

g) Sokal-Michener 20% trim

h) Sokal-Michener 30% trim

Figure S4: $\Delta\langle RMSD\rangle$ of six protein-peptide REMD systems between native structure calculated with $med_{c_0}$ to native structure calculated using other PRIME techniques. The scatter plot corresponds to the left y-axis and the bars correspond to the right y-axis. (a)-(h) uses the indicated similarity indices and trimming.
